# Microglia-derived extracellular vesicles attenuate acute a-synuclein induced astrocyte inflammation

**DOI:** 10.64898/2026.05.11.724371

**Authors:** Matthew P. Nelson, Daisy Dong, Kathleen A. Maguire-Zeiss

**Affiliations:** Georgetown University, School of Medicine, Department of Neuroscience, Washington, DC, United States; Georgetown University Interdisciplinary Program in Neuroscience, Washington, DC, United States; Georgetown University College of Arts & Sciences, Department of Biology, Washington, DC, United States

**Author notes:** **<u>Corresponding Author Email:</u>** Kathleen A. Maguire-Zeiss.

**Keywords:** extracellular vesicles, neuroinflammation, miRNA, microglia, cytokines, resveratrol

## Abstract

Aggregates of misfolded α-synuclein (Syn) and neuroinflammation are pathological features of Parkinson’s disease (PD). These, misfolded conformations of Syn promote cytokine and chemokine signaling in the surrounding microenvironment by triggering activation of glial cells through pattern recognition receptors. Microglia and astrocytes act as innate mediators of the neuroimmune response in the brain by regulating inflammatory signaling via paracrine and autocrine forms of cell communication. Extracellular vesicles (EVs) represent a form of glial cell to cell communication that can regulate the glial neuroimmune responses depending on the phenotype of the donor cell. Research has shown that the contents of EVs can be altered via pharmacologically altering the donor cell which offers a potential avenue for the regulation of inflammation. As such, we analyzed enriched mouse cortical primary astrocytes and characterized their response to Syn exposure in the absence and presence of microglia-derived EVs. Using trans-resveratrol, a naturally occurring polyphenol implicated for its anti-inflammatory properties, as our pharmacological agent to generate an anti-inflammatory microglial-derived EV phenotype we found that EVs derived from resveratrol-treated microglia decreased the production of proinflammatory molecules in enriched astrocytes exposed to Syn. Sequencing of EV miRNAs revealed two miRNAs (miR-5099 and miR-115) with significant up-regulation in resveratrol EVs compared to control EVs. Astrocytes transfected with corresponding miRNA mimics prior to Syn exposure showed a dramatic decrease in inflammatory biomarker production. These findings show that microglia-derived EVs and their specific miRNA cargo can attenuate Syn-directed inflammation in astrocytes and may serve as a novel therapeutic for proteinopathies like PD.

## Introduction

Parkinson’s disease (PD) is a neurodegenerative disorder characterized by the loss of dopaminergic neurons and the accumulation of misfolded α-synuclein (Syn)(Alfaidi et al., 2025; Calabresi et al., 2023; Chen et al., 2022; Vidović and Rikalovic, 2022). While neuronal dysfunction and death are central to PD pathology, neuroinflammation is markedly increased in the brains of PD patients throughout disease progression(Araújo et al., 2022). Using positron emission tomography [(PET), [^18^F]NOS PET], Doot et al. 2022 showed elevated oxidative stress in PD brains compared to age-matched controls, implicating microglia as the nidus of inflammation(Doot et al., 2022). Likewise, a meta-analysis of the PET tracer targeting TSPO, a transmembrane protein expressed on microglia, in PD found that TSPO levels were increased in PD brains compared to aged-matched controls(Alfaidi et al., 2025; Zhang and Gao, 2022). Taken together, the evidence for ongoing neuroinflammation in PD suggests a major role for glial cells in PD pathogenesis.

It is widely accepted that this neuroinflammation is heavily driven by the accumulation of fibrillar Syn which also exacerbates disease pathogenesis’(Isik et al., 2023; Khairnar et al., 2022; Tansey et al., 2022). Typically, the microglial innate immune response can be triggered by specific conformations of Syn through activation of pattern recognition receptors and phagocytosis pathways’(Caplan and Maguire-Zeiss, 2018; Chou et al., 2021; Filippini et al., 2019; Isik et al., 2023; Tansey et al., 2022). Microglia express toll-like receptors, that directly interact with Syn and induce the Nuclear Factor kappa B (NF-κB) transcription factor pathway leading to neuroinflammation(Béraud et al., 2011a; Béraud and Maguire-Zeiss, 2012; Daniele et al., 2015a; Maguire-Zeiss, 2008; Su et al., 2009a). While microglia are considered the innate immune cells of the brain, astrocytes are also important for this immune response(Zhao et al., 2024). In fact, here, we show that Syn activates astrocytes to express and release proinflammatory cytokines and chemokines. Others have also shown a central role for astrocytes in PD(Tansey et al., 2022)’(Chou et al., 2021)’(Zhao et al., 2024). In fact, cytokines released by pro-inflammatory astrocytes are linked to increases in PD severity and serve as critical early biomarkers for PD’(Mogi et al., 1994; Qu et al., 2023; Xiromerisiou et al., 2022). Overall, astrocytes and microglia play an important role in PD pathology; however, treatment options targeting glial activation remain limited (Al-Ghraiybah et al., 2022).

As such, in this study, we explored how crosstalk between microglia and astrocytes may act as a critical target for intervention in glial inflammation. It is known that glia and neurons ‘communicate’ at the quadpartite synapse and that glial cells can influence the phenotype of neighboring cells. In 2017, Drago et al. also showed that ATP could modify the extracellular vesicles released from microglia changing the microglial interaction with astrocytes’(Drago et al., 2017; Matejuk and Ransohoff, 2020; Squadrito et al., 2017). Extracellular vesicles (EVs) have a lipid bilayer membrane-bound structure that are up-taken by neighboring or distance cells facilitating cell communication via their heterogeneous cargo(Jeppesen et al., 2024)’(Yang et al., 2024). Therefore, EVs represent an efficient method whereby one cell can regulate the action of another cell(Goetzl et al., 2016; Paolicelli et al., 2019; Rothhammer et al., 2018). Previous literature has shown that cultured cells treated under specific conditions can produce EVs capable of regulating pro-inflammatory cascades in recipient cells’(Izquierdo-Altarejos et al., 2025; Lee et al., 2025; Madhu et al., 2024). The ability of EVs to influence another cell’s immunophenotype, may be due to their heterogeneous cargo containing varieties of different proteins, lipids, mRNAs, and miRNAs (Madhu et al., 2024). In particular, miRNAs, small noncoding RNAs that regulate mRNA expression and translation, play key roles in regulating inflammation (Li et al., 2020)’(Zhou et al., 2021).

In this study, we sought to drive microglia toward an anti-inflammatory phenotype using resveratrol (trans-3, 5, 4’-trihydroxystilbene), a naturally occurring polyphenol compound with anti-inflammatory properties, in order to produce a corresponding anti-inflammatory EV phenotype’(Miao et al., 2025; Sun et al., 2023; Tufekci et al., 2021). Resveratrol reduces the synthesis of pro-inflammatory mediators and induces anti-inflammatory proteins in microglia polarizing them towards an anti-inflammatory phenotype(Liu et al., 2022)’(“Resveratrol promoted the M2 polarization of microglia and reduced neuroinflammation after cerebral ischemia by inhibiting miR-155: International Journal of Neuroscience: Vol 130, No 8,” n.d.). First, we tested the hypothesis that resveratrol-treated microglia-derived EVs would alter the activation state of Syn-exposed astrocytes. Next, we interrogated whether the microglial EV cargo changed in response to resveratrol. We report a decrease in Syn-induced astrocyte activation following treatment with EVs derived from resveratrol-treated microglia. We next identified two miRNAs that mediate the decreased proinflammatory phenotype.

Taken together, this work suggests that anti-inflammatory EVs and specific miRNAs attenuate astrocyte activation driven by pathogenic synuclein and these miRNAs represent a new therapeutic approach for synucleinopathies.

## Methods

### Glia preparation

Primary glia cultures were derived from postnatal C57/Bl6 mouse cortices (P0-3) as previously described (Daniele et al., 2014). Cortices were isolated, homogenized, and cultured for at least 16 days in Microglia Culture Media (MCM; 1x Minimal Essential Medium Earle’s [MEM] supplemented with: 1 mM L-glutamine, 1 mM sodium pyruvate, 0.6% v/v D-(+)-glucose, 100 μg/ml Penicillin/Streptomycin (P/S), 4% v/v Fetal Bovine Serum (FBS), 6% v/v Horse Serum, and 0.1% Primocin. Flasks were then shaken for 3 h to isolate microglia, which were subsequently plated in Microglia Growth Media containing 5% v/v fetal bovine serum (FBS; MGM; Minimum Essential Medium Earle’s (MEM), supplemented with 1 mM sodium pyruvate, 0.6% (v/v) D-(+)-glucose, 1 mM L-glutamine, 100 μg/mL penicillin/streptomycin, and 5% v/v FBS)(Béraud et al., 2011a, 2011b; Béraud and Maguire-Zeiss, 2012; Daniele et al., 2015b). For RNA isolation, cells were plated at 8 × 10^5^ cells per well (6-well format). For immunocytochemistry, microglia were plated on sterile glass coverslips (12 mm; Deckglaser) at density of 4.0 × 10^4^ cells per well (24-well format). After microglia were removed from the mixed glia, enriched astrocytes were isolated by trypsinization, replated for 4-7 days, and grown until confluent. The enriched astrocytes were resuspended in MEM (supplemented with 10% FBS, 1 mM L-glutamine, and 100 μg/mL P/S) and plated in 24-well plates on glass cover slips at 4.0-5.0 × 10^4^ cells per well for immunocytochemistry or in 6-well plates at 5.0 × 10^5^ cells per well for mRNA analysis/protein assays. For all experiments, enriched astrocytes were serum-starved for 2 hours (MEM supplemented with 1 mM L-glutamine and 100 μg/mL P/S) prior to treatment.

### Resveratrol Preparation and Treatment

Trans-resveratrol (Sigma-Aldrich, Catalog # 1602105) was reconstituted in 100% ethanol (EtOH) at 40 mg/ml. Microglia were treated with 50 μM of resveratrol or an equal volume of EtOH vehicle in MGM containing 5% exosome depleted FBS (Gibco, Catalog # A2720803).

### Preparation and misfolding of α-synuclein

Human α-synuclein (Syn) was prepared as previously described(Daniele et al., 2015c; Giasson et al., 1999; Maguire-Zeiss et al., 2006). To misfold Syn, the lyophilized protein was resuspended to a final concentration of 0.1mg/mL in dPBS than sonicated in 1 second pulses for 2 minutes at 10% power using a 150E Sonic Dismembrator with 4C15 Probe. Cells were exposed to 1 μg/mL of Syn for all experiments.

### Immunocytochemistry

Following treatment, cells were washed with PBS for 5min prior to fixing in 4% paraformaldehyde/sucrose (w/v in PBS) for 20 min at room temperature. Cells were subsequently rinsed with PBS for 5min with gentle rotation and permeabilized for 5min in PBS containing 0.1% Triton X-100 with gentle rotation. For blocking, PBS containing 10% goat serum was applied for 1 h at room temperature with gentle rotation. Cells were incubated with primary antibodies for 24 h at 4°C in PBS containing 0.1% Triton X-100, and 1% goat serum with gentle rotation. Primary antibodies against GFAP (1:1000 Cell Signaling, Catalog # 36705), IBA-1 (1:1000 WAKO, Catalog # 019-19741), were used to identify astrocytes and microglia. Cells were washed 3 times for 10min with PBS + 0.1% Triton X-100 at room temperature with gentle rotation. At room temperature, cells were incubated with secondary antibodies for 1 hour in PBS containing 0.1% Triton x-100 and 1% goat serum. The secondary antibodies used were goat anti-rabbit IgG (H + L) highly cross-absorbed secondary antibody Fluor™ 488 (dilution 1:1000, Invitrogen, Catalog # A32731) and goat anti-rat IgG (H + L) highly cross-absorbed secondary Fluor™ Plus 594 (1:1000, Invitrogen, Catalog # A48264). Cells were washed again for 3 × 10min with PBS + 0.1% Triton X-100 at room temperature with gentle rotation. Coverslips containing cells were mounted using Vectashield Vibrance antifade mounting medium with DAPI (Vector, Catalog # H-1800-10).

### Microscopy and Analysis

Researchers were blinded during image acquisition and analyses. Cells were imaged at 10x, 20x, and 40x magnification using Zeiss Axio Imager Z2 with Axiocam 705-Mono scope. Images were captured using capture software Zeiss Zen 3.9. Four images, equal distance from each other, were acquired per coverslip. Each image contained at least 20 cells which were identified by DAPI nuclei staining.

### Characterization of mixed glia

The specific markers GFAP and IBA1, were used to measure the density of astrocytes and microglia respectively. Nuclei staining with DAPI was used to identify regions with more than 20 cells. Images were taken at 20x magnification. For each image, the total number of DAPI cells was calculated using the Image-J analysis particle plug-in. IBA-1 and GFAP expressing cells were then counted manually. The percentage of IBA-1 and GFAP positive cells was calculated relative to the total number of DAPI stained nuclei. Four images from 5 coverslips were analyzed for a total of 20 images per group.

### Quantitative Real-Time PCR

Total RNA was extracted from enriched astrocyte cultures using the RNeasy Mini Kit (Qiagen, Catalog # 74104) and transcribed into cDNA using the High-Capacity RNA-to-cDNA Kit (Applied Biosystems, Catalog # 4368814). Gene expression was quantified using qRT-PCR in a 384 well format in an ABI Prism 7900HT Sequence Detection System (Life Technologies). Data were analyzed using the relative delta-delta Ct method of quantification. 18S and GAPDH were used as reference genes and fold changes calculated using the ΔΔCt method, with statistical analyses performed on ΔCt values. Gene expression changes are represented as fold change (2^-ΔΔCt^). All measurements were performed with at least three biological replicates, three technical replicates, and three experimental replicates.

### Exosome isolation from microglia

Microglia were cultured in Dulbecco’s Modified Eagle’s Medium with 5% exosome depleted FBS (Gibco, Catalog # A2720803). Microglia were treated with 50 µM of resveratrol (Sigma-Aldrich, Catalog # R5010) or an equal volume of vehicle (EtOH) for 24 hr and extracellular vesicles from each condition isolated as described here: microglia supernatants were collected and centrifuged at 2000 g for 30 min, supernatant was added to Exosome Isolation Kit (Invitrogen, Catalog # 4478359) followed by centrifugation at 10k g for 1 h according to manufacturer’s specifications. The resulting extracellular vesicles (EVs) derived from vehicle-treated microglia (VEVs) and resveratrol-treated microglia (REVs) were resuspended in dPBS, RLT Buffer (Qiagen, Catalog #74106), or TRIzol reagent (Invitrogen, Catalog # 15596026) and subsequently used for enriched astrocyte treatment or analyzed. EVs were stored at -80 °C until use.

### Electron Microscopy

EVs derived from resveratrol and vehicle treated microglia were resuspended dPBS and 5 µL of the EV suspensions were applied to formvar/carbon coated 200 mesh copper grids (Electron Microscopy Sciences) that were glow discharged (20 mA for 60 seconds) just prior to use. EVs were negatively stained with 1% uranyl acetate (Electron Microscopy Sciences) for 1 minute and air dried before imaging. A FEI Talos F200X transmission electron microscope operated at 200 kV was used to acquire images. Sample preparation and imaging was performed by the George Washington University Nanofabrication and Imaging Center.

### Nanoparticle Tracking Analysis

Microglia-derived EVs were subjected to nanoparticle tracking analysis (NTA) using the Nanosight NS3000. EVs were diluted (1:10) in UltraPure Distilled Water (Invitrogen, Catalog #10977-015) and analyzed. Particle-size distribution and concentration were measured by capturing the particles undergoing Brownian movement and tracking the trajectories using laser light for particle quantification. NTA 3.0 software was used for all analyses. Each sample was analyzed 5 times for 60 sec. Mean size and concentration of EVs/ml were calculated for five replicate experiments.

### ELISAs

The levels of TNF-α, IL-6, CCL2, and IL-1β (R&D systems) in the enriched astrocyte conditioned media were quantified using ELISA kits and a microplate reader (Molecular Devices, Spectra Max iD3), according to the manufacturer’s instructions.

### Western Blots

Protein concentrations of EVs and total cell lysates were determined using a colorimetric protein assay (Bio-Rad, Catalog #5000001). Protein samples were diluted in Laemmli sample buffer containing beta-mercaptoethanol (5%). Samples were then heated for 10 minutes at 80 ° c. Proteins were separated by denaturing polyacrylamide gel electrophoresis and transferred to PVDF membranes. Proteins were stained with Ponceau S and imaged, followed by blocking for 1 hour at room temperature in TBST (20 mM Tris-HCl, pH 7.5, 150 mM NaCl, 0.1% (v/v) Tween 20) and 5% (w/v) non-fat dried milk (NFDM) with gentle rocking. Membranes were then incubated overnight at 4°C in TBST/5%NFDM with primary antibodies (TSG101 1:1000 Abcam, Catalog # ab30871; CD63 1:1000 Abcam, Catalog # Ab217345). The membranes were washed and then incubated with horse radish peroxidase-conjugated goat anti-rabbit (1:2000 Invitrogen, Catalog # 31460) for 1 hour at room temperature with gentle rocking. SuperSignal™ West Pico PLUS Chemiluminescent Substrate (Thermofisher, Catalog # 34579) was applied to identify the antigen-antibody complexes. Membranes were imaged using GE Amersham 600 RGB and Azure 300 Chemiluminescent Imaging system.

### miRNA Sequencing and Analysis

Next generation miRNA sequencing was performed by Azenta Genewiz and the raw FASTQ files were stored for miRNA sequencing analysis. RStudio was used to perform the analysis. Differentially expressed genes were identified using DESEQ2 package(Love et al., 2014). All data received from DESEQUE analysis was exported to excel files for downstream analysis and visualization. The *P* values of multiple testing were adjusted and a cutoff of p_adj_<0.05 and Log2(Fold change) >0.5. Of those miRNAs, potential targets for miRNA-5099 were identified using miRDB. Those targets were used as the criteria genes used for ShinyGo 0.85.1 and mapped to the mouse(“Ensembl 2025 | Nucleic Acids Research | Oxford Academic,” n.d.; “KEGG: integrating viruses and cellular organisms | Nucleic Acids Research | Oxford Academic,” n.d.; “Pathview: an R/Bioconductor package for pathway-based data integration and visualization | Bioinformatics | Oxford Academic,” n.d.; Ge et al., 2020; Love et al., 2014). GraphPad was used for data visualization. The size of the circle references the number of genes (nGenes) from the geneset in the pathway, the color represents the False Detection Rate (FDR). The y axis is the labeled pathway associated with the genes. The x axis refers to the Fold Enrichment (FE) which refers to the nGene divided by the total number of Pathway Genes (pGenes).

### miRNA Transfection

Lipofectamine™ RNAiMAX Transfection Reagent (Invitrogen, Catalog # 13778100) was used according to manufacturing protocols for 24-well plate format. Enriched astrocytes were plated (4×10^4^ cells/well; 24-well format) and transfected 48 hours after plating. Cells were transfected 24 hours prior to additional treatments. miRNA mimics for mmu-miR-5099 (MED Chem express, Catalog# HY-R03245), mmu-miR-720 (Thermofisher, Catalog # 4427975) which has the same mature sequence as target Novel-miRNA-115 identified though miRNA sequencing, and scrambled miRNA (MCE, Catalog # HY-R04602 Lot#825593) were utilized for the transfections. Lipofectamine alone was used as a transfection control. Astrocytes were then treated with 1 μg/ml of Syn or vehicle for 24 hours. Conditioned media was stored at -80 °C prior to analysis via ELISAs.

## Results

### Pathogenic α-synuclein induces activation of cultured astrocytes

Using enriched astrocytes derived from mouse cortices, we first established the glial composition the cultures (**Fig. 1a**). We next quantified inflammatory cytokine expression following exposure to pathogenic vehicle or α-synuclein (Syn; **Fig. 1b & c**). Gene expression of IL-6, CCL2, TNF-α and IL-1β were all significantly increased in enriched astrocytes following Syn exposure compared to vehicle conditions (**Fig. 1b**). Subsequent analysis of the conditioned media from treated cells showed that protein expression levels for the corresponding inflammatory factors were also increased (**Fig. 1c**).

**Fig. 1.**
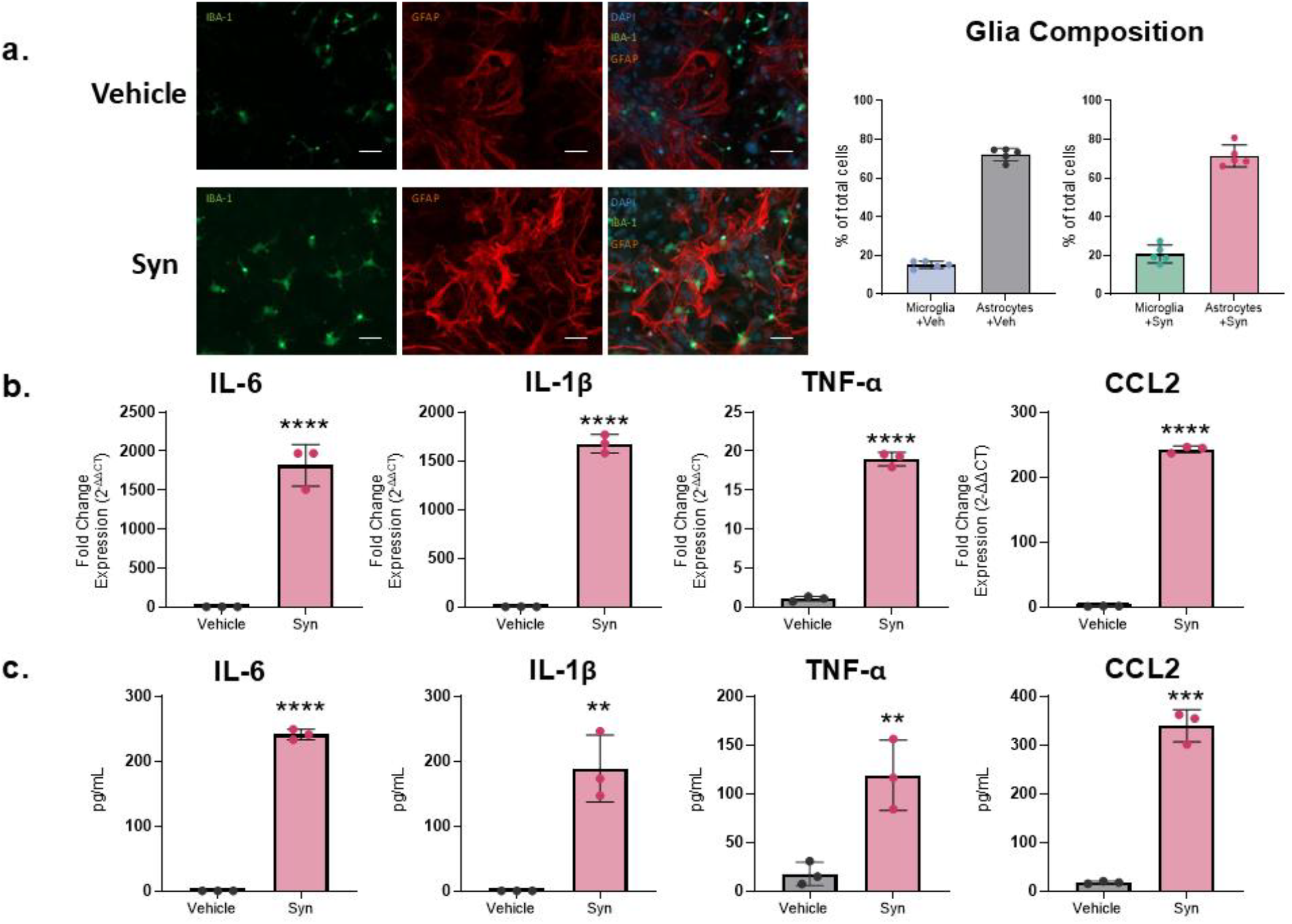
Syn induces glial activation: **a)** Representative images of IBA1+ microglia and GFAP+ astrocytes in mixed glia cell cultures derived from mouse cortical tissue. The composition of the mixed glia was ∼15% microglia and ∼72% astrocytes (referred to as enriched astrocytes; nuclei stained with DAPI; 10x magnification with 100 µm scale bar **b)** qRT-PCR analyses of proinflammatory gene expression in enriched astrocytes 24 hours after vehicle or Syn treatment. **c)** Quantification of proinflammatory protein levels in enriched astrocyte conditioned media 24 hours after vehicle or Syn treatment via ELISAs. All analyses were performed using two-tailed student’s t-tests *P* = ** <0.01, *** <0.001, **** < 0.0001, mean ± SEM; n=3; with 3 technical replicates per condition and 3 experimental repeats

### EVs derived from resveratrol-treated microglia attenuate Syn-activation of astrocytes

It is well known that glial cells communicate with each other and neurons to maintain homeostasis in the brain. Extracellular vesicles are lipid-bound particles secreted by all cells and can contain proteins, microRNAs, mRNAs, DNA, and lipids facilitating cell-cell communications. Because Syn increased the inflammatory signature of enriched astrocytes and since microglia and astrocytes both play an important role in the brain’s response to neuroinflammation, we next asked whether extracellular vesicles derived from microglia pretreated with an anti-inflammatory drug could change the inflammatory profile of Syn-exposed astrocytes. To that end, primary microglia were treated with the anti-inflammatory drug, resveratrol (50 µM). Resveratrol alone did not alter the expression of IBA1 in resting microglia (**Fig. 2a**). EVs were then isolated from vehicle-treated and resveratrol-treated microglia (VEVs and REVs, respectively). Both types of EVs showed the characteristic morphology and size of EVs following transmission electron microscopy (TEM) imaging (**Fig. 2b**). Western blot analysis confirmed the enrichment of extracellular vesicle markers CD63 and TSG101 in both VEVs and REVs with a relative decrease in whole cell microglia lysates (**Fig. 2c)**. The average particle size was not significantly different between treatment groups nor was the number of microglia-derived EVs using nanoparticle tracking analysis (NTA) (**Fig. 2d**). As shown in the distribution curve, both VEVs and REVs exhibited a similar size distribution. Together this data shows that EVs derived from vehicle and resveratrol treated microglia exhibit similar morphological characteristics and distributions of small and large EVs(Welsh et al., 2024).

**Fig. 2.**
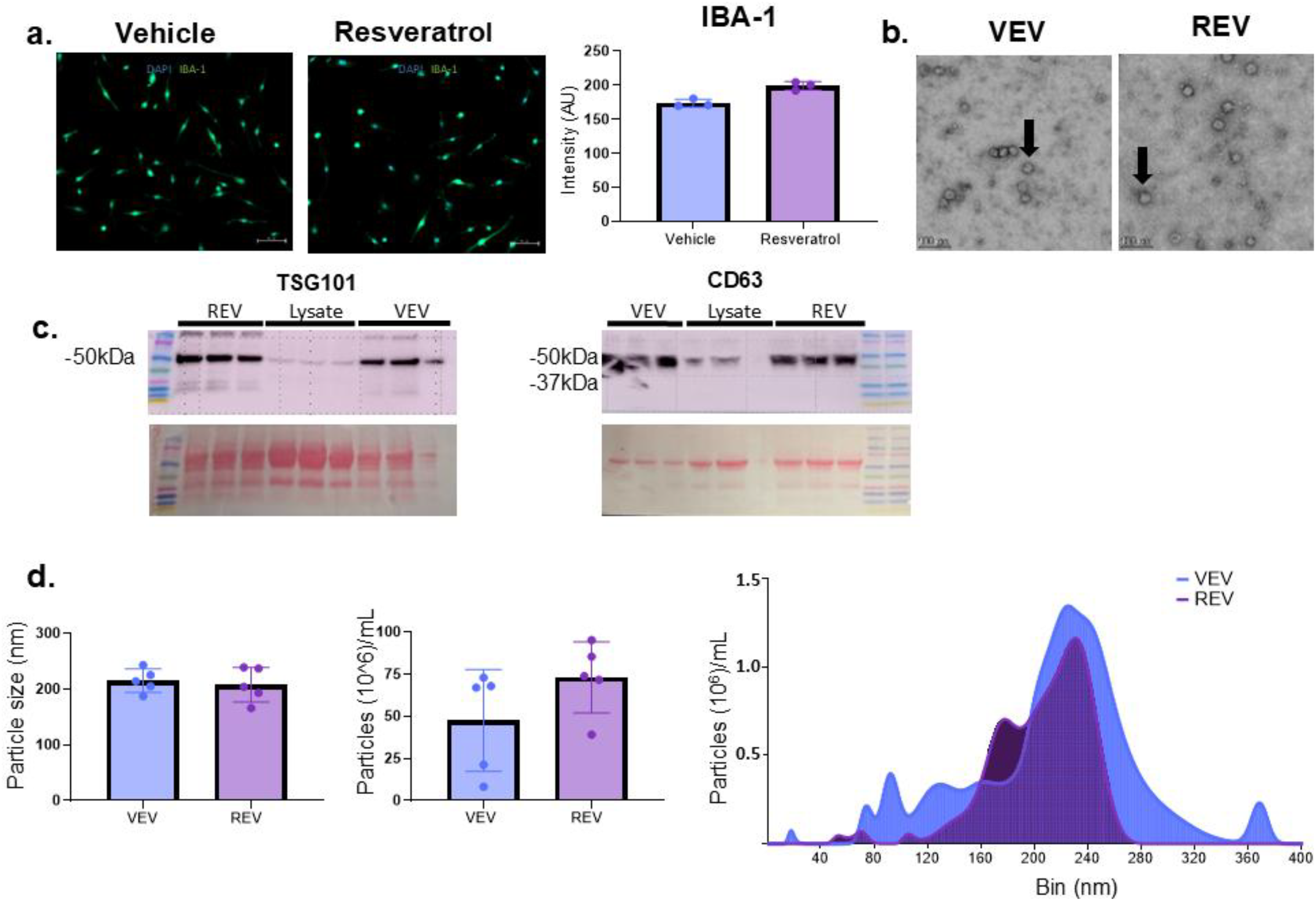
Characterization of resveratrol treated microglia and extracellular vesicles (EVs): **a)** Representative images and analysis of IBA1+ microglia 24 hours after vehicle or resveratrol treatment; nuclei stained with DAPI; 20x magnification with 50 µm scale bar n=5 coverslips per condition with 4 images per coverslip = 20 images per condition; the intensity (AU; arbitrary units) of IBA1+ staining was not significantly different between resveratrol and vehicle treated microglia **b)** Representative Transmission Electron Microscopy (TEM) images of EVs obtained from vehicle (VEV) and resveratrol (REV) treated microglia. Scale bar, 100 µm; black arrows point to representative particles with morphology meeting the criteria of EVs **c)** Representative western blots of EV proteins, TSG101 and CD63, in REVs, VEVs, and whole cell lysates; membranes were stained with Ponceau S for total protein loading; n=3 technical replicates **d)** Quantification of extracellular vesicle size, concentration, and distribution performed via Nanoparticle Tracking Analysis (NTA); n=5 biological replicates; all analyses were performed using two-tailed student’s t-tests; there were no significant differences in particle size, concentration or distribution

To assess the impact of REVs on Syn-induced inflammation, we treated enriched astrocyte cultures with VEVs, REVs, or the drug, resveratrol (Res), while stimulating activation using Syn exposure. Twenty-four hours later we interrogated the morphofunctional changes of these astrocytes. First, we note that there are no discernable morphological differences between enriched astrocytes treated with REVs and Syn and those treated with Syn alone (**Fig. 3a**). However, quantification of proinflammatory gene expression (IL-6, IL-1β, TNF-α, and CCL2) revealed a significant decrease in expression of these mRNAs in the REV treatment groups. Interestingly, the VEVs also decreased the expression of a subset of the mRNAs (TNF-α and IL-1β; **Fig. 3b**). Protein expression of IL-6, IL-1β, TNF-α, and CCL2 showed a signification decrease in the REV treated astrocyte-conditioned media compared to Syn exposure alone (**Fig. 3c**). VEV treatment also decreased the expression of Il-6, IL-1β, and CCL2 in the astrocyte-conditioned media. Resveratrol alone only decreased the release of CCL2 by 30% but REV treatment significantly decreased gene expression compared to resveratrol. In all cases, REV treatment had a significant effect on Syn**-**induced inflammation in enriched astrocytes.

**Fig. 3.**
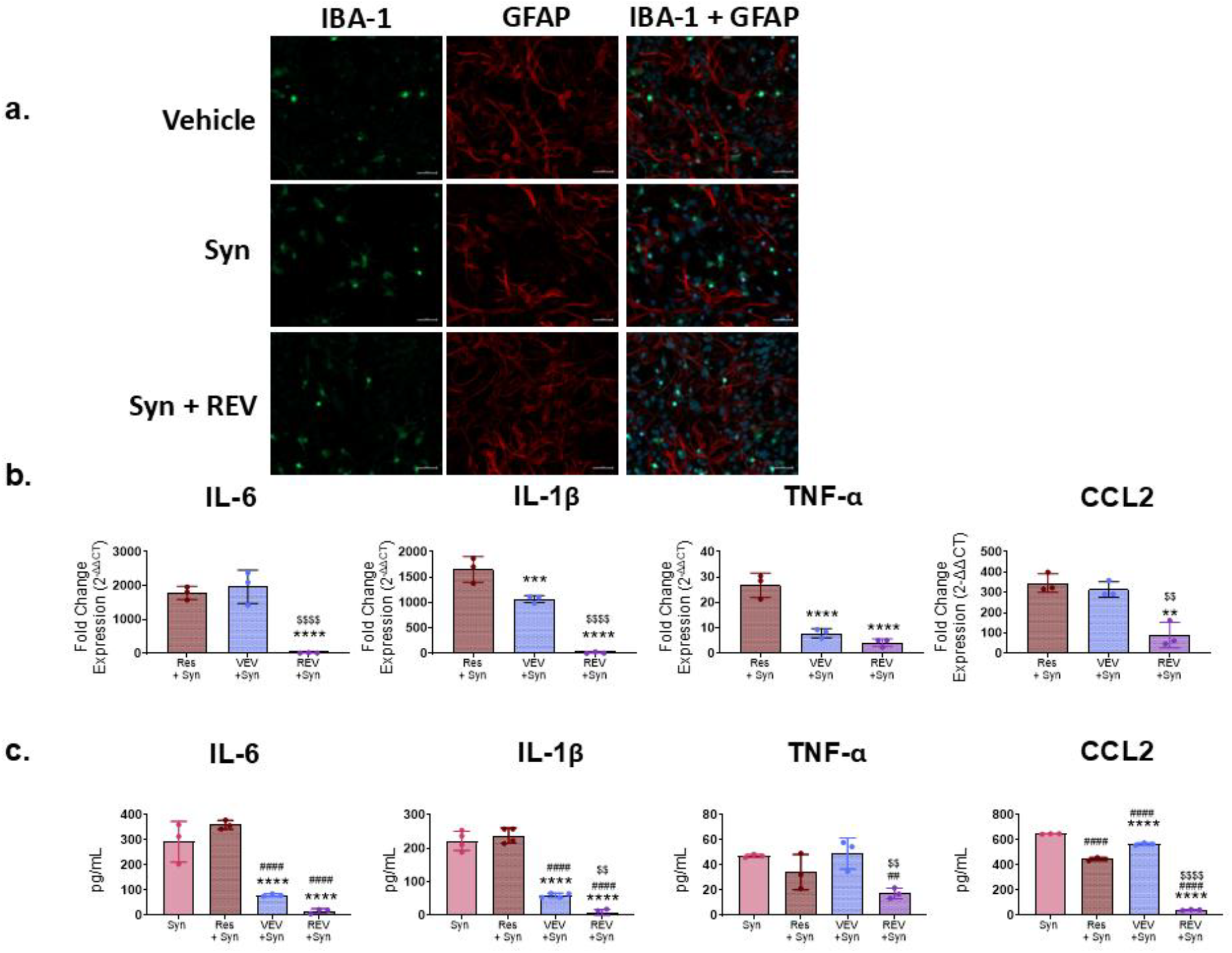
Extracellular vesicles attenuate Syn-induced Inflammation: **a)** Representative images of IBA1+ microglia and GFAP+ astrocytes in enriched astrocyte cultures derived from mouse cortices treated with vehicle (PBS), Syn, or Syn + REVs; nuclei stained with DAPI, 10x magnification; scale bar 100 µm **b)** qRT-PCR analyses of proinflammatory gene expression in enriched astrocytes 24 hours after exposure to Syn plus resveratrol, VEVs or REVs. **c)** Quantification of proinflammatory proteins in enriched astrocyte conditioned media 24 hours after exposure to Syn, Syn plus resveratrol, VEVs or REVs via ELISAs; all analyses were performed using one-way ANOVA; *P* =**^$$##^ <0.01, ***^$$$##^ <0.001, ****^$$$###^ < 0.0001; *=relative to RES + Syn; ^$^ =relative to VEV; ^#^=relative to Syn; mean ± SEM; n=3; with 3 technical replicates per condition; this experiment repeated 3 times with similar results

### Resveratrol induces changes in the miRNA profile of microglia-derived EVs

Since microglia-derived REVs significantly and consistently decreased the Syn-induced proinflammatory profile of the astrocytes, we next sought to identify the miRNA profile of the EVs. High throughput miRNA sequencing of microglia-derived VEVs and REVs revealed 837 miRNA targets. For this analysis, miRNAs with an adjusted p-value (Padj) >0.5 and log base 2-fold change [log2(FC)] >9.0 were selected for further analysis. Eleven miRNAs were found to be significantly upregulated and 13 were found to be downregulated following resveratrol treatment (**Fig. 4a**). Of the upregulated miRNAs, we identified 2 highly increased miRNAs from REVs: miR-5099 and miR-115. To better understand the possible pathways affected by resveratrol, we performed a KEGG analysis of miR-5099 targets **(Fig. 4b)**, using the miRNA target prediction database(Chen and Wang, 2020; “Prediction of functional microRNA targets by integrative modeling of microRNA binding and target expression data | Genome Biology | Springer Nature Link,” n.d.). We included only targets with a target prediction value >80. The greatest enrichment was in the proteosome category, as well as other pathways related to PD and neurodegeneration suggesting that miR-5099 may be a potential therapeutic agent for proteinopathies.

**Fig. 4.**
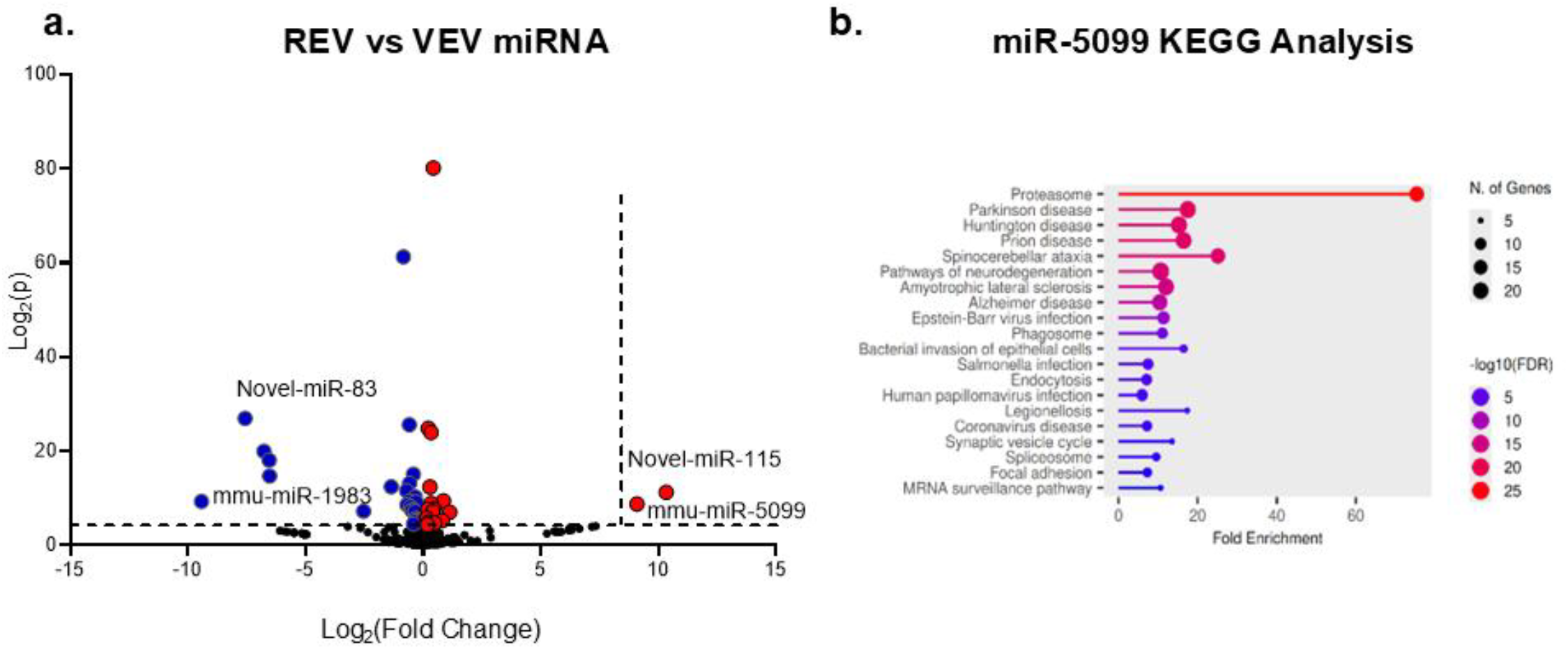
Resveratrol induces changes in the miRNA profile of microglia-derived extracellular vesicles: **a)** Volcano plot representation of miRNA expression differences between microglia-derived VEVs and REVs following miRNA sequencing analysis; Upregulated genes (red) and down regulated genes (blue) with Log_2_foldchange>4 and Log_2_p value>4 for significance **b)** KEGG pathway analysis of miR-5099 associated target proteins determined via miRNA target analysis

### miRNA mimics attenuate Syn-induced activation of enriched astrocyte

Since miR-5009 and miR-115 showed between a 8- and 10-fold increase in microglia-derived REVs compared to VEVs, we transfected enriched astrocytes with corresponding miRNA mimics or scrambled controls. Following transfection with each mimic alone and in combination the enriched astrocytes were activated with Syn for 24 hours. Cytokine protein expression was subsequently quantified in condition media from these cells. As shown in (**Fig. 5**), transfection with the combination of miR-5099 and -115 decreased the Syn-induced proinflammatory profile for all targets compared with the scrambled control. Each mimic alone had less of an effect on IL-1β, TNF-α, and CCL2 expression compared with the combined mimic treatment.

**Fig. 5.**
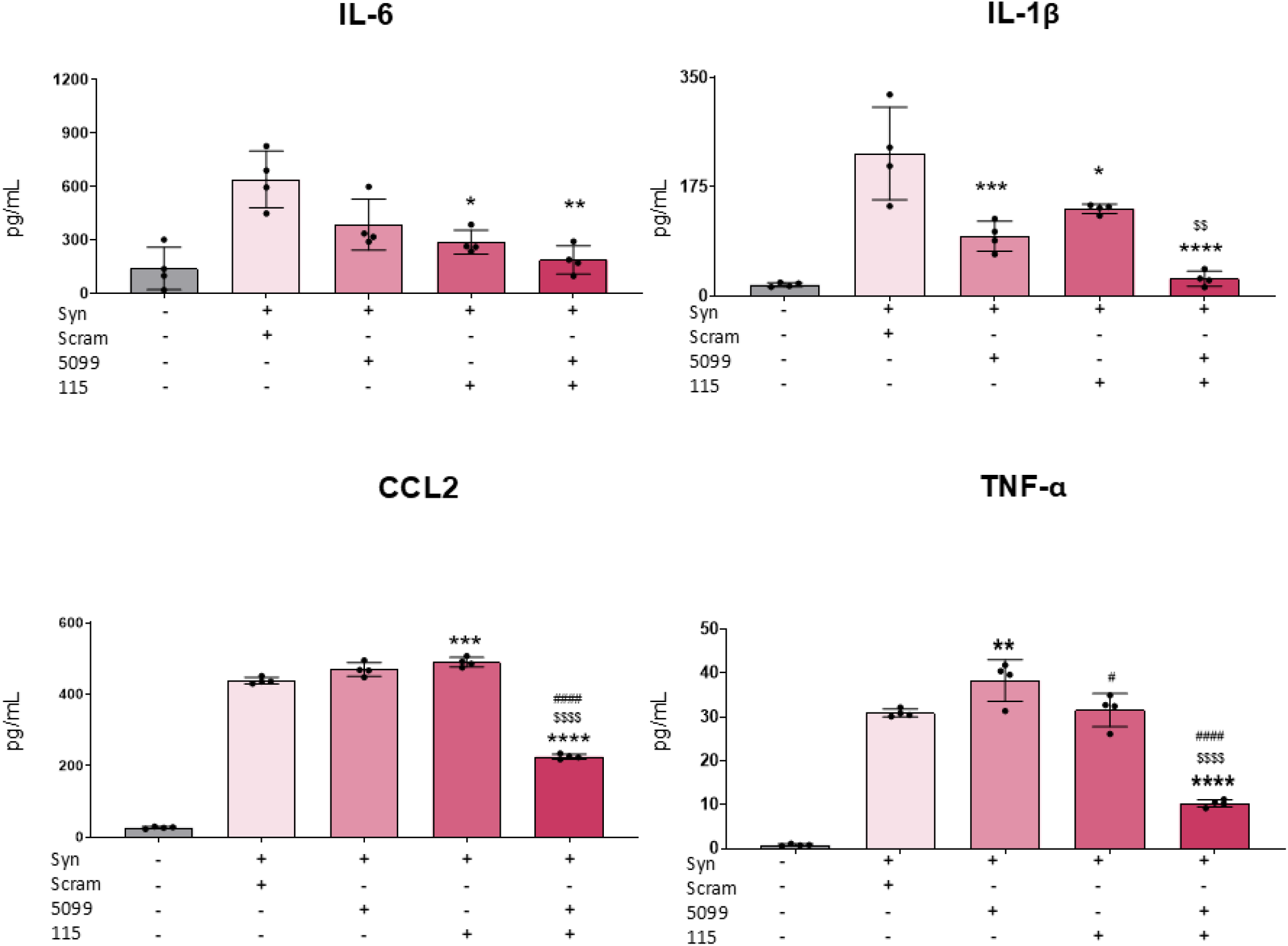
miRNA mimics attenuate Syn-induced activation of enriched astrocytes: **a)** Quantification of proinflammatory proteins from condition media of enriched astrocytes transfected with miRNA mimics or scrambled (Scram) nucleotide controls, 24 hours after exposure to Syn or vehicle via ELISAs; all analyses were performed using one-way ANOVA; *P* =*^$#^=0.05, *P* =**^$$##^ <0.01, ***^$$$##^ <0.001, ****^$$$###^ < 0.0001, *=relative to Syn + Scram; ^$^ =relative to Syn + 115 mimic ^#^=relative to Syn + 5099; mean ± SEM; n=3; with 3 technical replicates per condition; this experiment repeated 3 times with similar results

## Discussion

Our findings provide insight into a novel mechanism for regulating astrocyte inflammation. Astrocytes play a critical role, in Parkinson’s disease (PD) neurotoxicity, by the release of inflammatory cytokines and in the phagocytosis of Syn(Filippini et al., 2019; Rostami et al., 2021). Therefore, astrocyte modulation may regulate PD inflammation severity and neurodegeneration^56–58^. Utilizing an *in vitro* model of synuclein-induced inflammation, we identified a pharmacological intervention that shifted microglia to produce extracellular vesicles (EVs) with greater efficacy in reducing inflammation in astrocytes. The primary mouse glial cell model allows us to study microglia-astrocyte communication and providing a means of isolating each cell type and characterizing them accordingly. First, we confirmed that enriched astrocytes exposed to Syn adopt a proinflammatory phenotype expressing and releasing higher levels of IL-6, IL-1β, TNF-α, and CCL2 compared with unstimulated astrocytes. These findings are not surprising given that astrocytes express toll-like receptors (TLRs) and previous research from our lab and others has shown that Syn is a TLR ligand activating the NF-κB transcription factor pathway to increase expression of these molecules (Béraud et al., 2011a; Daniele et al., 2015b; Fellner et al., 2013; Isik et al., 2023; Li et al., 2021; Su et al., 2009b).

Previous studies demonstrated that resveratrol induces an anti-inflammatory polarization in microglia, reducing oxidative stress, and preventing cytokine release via modulation of NF-κB translocation making it a potentially effective intervention. However, while resveratrol has good solubility and can cross the blood brain barrier it has limited bioavailability and novel methods to improve efficacy are needed(Chinraj and Raman, 2022; Liu et al., 2022; Sun et al., 2023; Tufekci et al., 2021)’(Kogut et al., 2026). Furthermore, in our hands, there was no significant anti-inflammatory effect of resveratrol alone on enriched astrocytes cotreated with synuclein. These findings led us to consider manipulating the cell-cell communication between microglia and astrocytes as an alternative method to attenuate inflammation. We reasoned that because cell-cell communication is an essential part of ongoing neuroinflammation as well as the brain’s drive to homeostasis, manipulating that communication to alleviate inflammation may serve as a potential new therapeutic approach. As with all cells, this paracrine communication requires signals to be transduced by the receiving cell into information that allows for a specific function. Here, we forced the phenotype of microglia towards an anti-inflammatory signature by treating these microglia cells with resveratrol. We next isolated and characterized the extracellular vesicles (EVs) and tested whether Syn-directed astrocyte activation was attenuated.

Our study provides evidence that astrocyte activation is attenuated by microglial EVs. We found that naive microglia-derived EVs dampened the inflammatory profile of astrocytes suggesting an innate ability of microglia to regulate the astrocyte response to Syn. However, the EVs from resveratrol-treated microglia had a more robust effect on Syn-directed astrocyte activation with a significant reduction in the expression and release of all the inflammatory molecules quantified (IL-6, IL-1β, TNF-α, and CCL2).

This reduction in proinflammatory molecules regulated by NF-κB led us to examine the EV contents, since EVs are secreted from all cells and are important for intercellular communication their contents can change significantly. Furthermore, EVs are well-known to transport a variety of different materials including miRNAs, lipids, and proteins(Jeppesen et al., 2024; Paolicelli et al., 2019). We focused this study on miRNAs, because they modulate gene expression, are highly conserved, and can exhibit tissue and cell specificity(Bartel, 2004; Mori et al., 2019). In addition, the expression of noncoding small RNAs are implicated in many disorders including Parkinson’s disease(Elangovan et al., 2025; Goh et al., 2019; Nair and Ge, 2016; Shaheen et al., 2024). For example, analyses of postmortem Parkinson’s disease tissue revealed altered miRNA profiles in both the striatum and substantia nigra(Briggs et al., 2015; Nair and Ge, 2016). Many pathways are implicated in these studies including oxidative stress, protein degradation, synaptic plasticity and inflammation.

In our study, sequencing of microglia-derived EVs revealed two miRNAs with a greater than 8-fold change in expression following resveratrol treatment, miRs -5099 and -115. Subsequent transfection of astrocytes with a combination of miR-5099 and miR-115 mimics decreased Syn-directed astrocyte inflammation, suggesting a novel therapeutic target for neuroinflammation. Previous studies have investigated these miRNAs independently. In a model of obesity, Tang et al altered the adipocyte-derived miRNA profile using an anti-inflammatory drug and showed an upregulation of miR-5099 which then attenuated NF-κB mediated inflammation in macrophages(Tang et al., 2026). While miR-115, a novel miRNA, is implicated in tumor cell growth with very little known about its cellular targets(Xu and Shi, 2019). However, our study noted a particularly significant anti-inflammatory effect when transfected together suggesting that EVs produce miRNAs with synergistic properties.

In conclusion, we found that microglia-derived EVs can alleviate pro-inflammatory signaling in astrocytes activated by synuclein. We further identified two miRNAs, miR-5099 and miR-115, upregulated in resveratrol-treated microglia EVs that when transfected into astrocytes significantly decreased the release of pro-inflammatory cytokines. Since ongoing expression of pro-inflammatory cytokines is detrimental to brain health and exacerbates neurodegeneration(Adamu et al., 2024; Barcia et al., 2011; Filippini et al., 2019; Hennessy et al., 2017; Zhang et al., 2023), identifying druggable targets for therapy is essential. We suggest that the findings of this study provide a potential new target for dampening synuclein-induced glial activation and subsequent neuroinflammation.

## List of Abbreviations

Syn: α-synuclein
EVs: Extracellular Vesicles
IL-6: Interleukin-6
CCL2: chemokine (C-C motif) ligand 2
TNF-α: Tumor necrosis factor
Il-1β: Interleukin-1 beta
NF-κB: Nuclear factor kappa-light-chain-enhancer of activated B-cells
TLR: Toll-like receptors
PD: Parkinson’s disease

## Acknowledgements

We acknowledge and thank Dr. Amrita Cheema and Sunain Deol for their assistance with particle analysis at the Georgetown University center for Metabolomic Studies; as well as, the George Washington University Nanofabrication and Imaging Center, the Georgetown University Mass Spectrometry and Analytical Shared Resources, and the Georgetown University Division of Comparative Medicine veterinarians and veterinary technicians for the care of the mice.

## Author contributions

MN and KM-Z designed the experiments. MN performed the experiments and wrote the first draft of the manuscript. DD assisted with microglia immunostaining and image taking. MN and KM-Z analyzed data, wrote sections the manuscript, and contributed to manuscript revision. All authors read and approved of the submitted version.

## Ethics approval

All procedures and experiments were approved by Georgetown University’s Institutional Animal Care and Use Committee and followed the United States’ National Institute of Health ethical guidelines. Animals were housed in Georgetown University Division of Comparative Medicine facilities with ad libitum water and food in a 12H light/dark cycle.

